# Sediment-associated microbial community profiling: sample pre-processing through sequential membrane filtration for 16s rDNA amplicon sequencing

**DOI:** 10.1101/2020.10.21.348342

**Authors:** Joeselle M. Serrana, Kozo Watanabe

## Abstract

Sequential membrane filtration as a pre-processing step for the isolation of microorganisms could provide good quality and integrity DNA that can be preserved and kept at ambient temperatures before community profiling through culture-independent molecular techniques, e.g., 16s rDNA amplicon sequencing. Here, we assessed the impact of pre-processing sediment samples by sequential membrane filtration (from 10, 5 to 0.22 μm pore size membrane filters) for 16s rDNA-based community profiling of sediment-associated microorganisms. Specifically, we examined if there would be method-driven differences between non- and pre-processed sediment samples regarding the quality and quantity of extracted DNA, PCR amplicon, resulting high-throughput sequencing reads, microbial diversity, and community composition. We found no significant difference in the quality and quantity of extracted DNA and PCR amplicons between the two methods. Although we found a significant difference in raw and quality-filtered reads, read abundance after bioinformatics processing (i.e., denoising and the chimeric-read filtering steps) were not significantly different. These results suggest that read abundance after these read processing steps were not influenced by sediment processing or lack thereof. Although the non- and pre-processed sediment samples had more unique than shared amplicon sequence variants (ASVs), we report that their shared ASVs accounted for 74% of both methods’ absolute read abundance. More so at the genus level, the final collection filter identified most of the genera (95% of the reads) captured from the non-processed samples, with a total of 51 false-negative (2%) and 59 false-positive genera (3%). Accordingly, the diversity estimates and community composition were not significantly different between the non- and pre-processed samples. We demonstrate that while there were differences in shared and unique taxa, both methods revealed comparable microbial diversity and community composition. We also suggest the inclusion of sequential filters (i.e., pre- and mid-filters) in the community profiling, given the additional taxa not detected from the non-processed and the final collection filter. Our observations highlight the feasibility of pre-processing sediment samples for community analysis and the need to further assess sampling strategies to help conceptualize appropriate study designs for sediment-associated microbial community profiling.

## INTRODUCTION

Microorganisms have long been recognized as useful bioindicators for biomonitoring and ecological assessment of freshwater ecosystems (Payne, 2013; Amleida et al., 2014; Pawlowski et al., 2016). Recent studies took advantage of high-throughput sequencing (HTS) to characterize freshwater sediment-associated microorganisms for impact assessment of anthropogenic activities and environmental factors on diversity and composition and their functions (e.g., Stern et al., 2017, Liao et al., 2019). In particular, 16s rDNA amplicon sequencing is a relatively faster and cheaper approach providing substantially higher taxonomic resolution (Singer et al., 2016), with the capability of detecting unculturable, rare, and novel microorganisms (Browne et al., 2016) in comparison to the conventional strategies, e.g., culture-dependent methods (Fransoza et al., 2015) for microbial community profiling.

Most studies would directly extract microbial DNA from sediment samples, amplify a target hypervariable region of the 16s rDNA gene through polymerase chain reaction (PCR), process for amplicon library construction, and sequence on a high-throughput platform (e.g., Illumina-based technologies). One major challenge with such an approach would be the isolation and capture of good quality and quantity DNA from sediment samples (Harnpicharnchai et al., 2007; Solomon et al., 2016), which mostly contain impurities that inhibit amplification through PCR (Albers et al., 2013). Various commercial extraction kits are available for the rapid processing of environmental samples tailored to yield abundant and high-quality DNA minimizing the effects of enzyme inhibitors, e.g., humic acid, polysaccharides, metals, etc. that must be removed before amplification with the help of proprietary chemicals (Kosch and Summer, 2013; Ni et al., 2016; Lear et al., 2018). However, most of these kits commonly rely on DNA-binding steps via silica spin columns for DNA purification and concentration. This procedure possibly results in DNA loss due to the competitive column-binding of organic matter (Lloyd, MacGregor, and Teske, 2010) that has been reported to selectively retain high molecular-weight DNA fragments (Rohland et al., 2018). Furthermore, collected sediments and other organic matter usually result in a large sample volume that requires proper processing so that DNA representing the whole community can be extracted, similar to environmental DNA samples (Aylagas et al., 2016).

Pre-processing sediment samples by multi-level or sequential membrane filtration have been reported to efficiently isolate high-quality DNA while reducing inhibitory enzyme compounds (Solomon et al., 2016; Kachiprath et al., 2017; Mathai et al., 2019; Sakami, 2019). Sequential filtration has been used to concentrate microbial biomass and assess communities based on size fractions using filter membranes with different pore sizes (Padilla et al., 2015; Bae, Lyons, and Onstad, 2019). A pre-filter of larger pore size (1.0 to 30 μm) and a collection filter of smaller size (0.22 μm) are commonly used in-line series of filters (Stewart et al., 2012; Liu et al., 2017) to efficiently capture viruses, bacteria, and parasites based on size exclusion (Hill et al., 2007). DNA is then extracted from the final collection filter to separate targeted microorganisms from the comparatively larger eukaryotic cells (e.g., Smith et al., 2017) or to remove large particle-associated microbes from the free-living fraction (e.g., Teeling et al., 2012; Smith et al., 2013; Orsi et al., 2015; Padilla et al., 2015; Schultz et al., 2020).

Previous studies have characterized and compared the microbial community structure of various collection strategies against *in situ* or on-site filtration of particle or sediment collected samples, mainly from marine environments (e.g., Puigcorbé et al., 2020; Torres-Beltrán et al., 2019). On-site filtration keeps the sampled microbial communities *in situ* conditions while reducing the time between collection and storage (Puigcorbé et al., 2020). The microorganisms from environmental samples should be inactivated right after collection without significant damage to their DNA (Song et al., 2016). Managing this time is critical to prevent bacterial overgrowth or taxonomically biased DNA damage and degradation (Hugerth and Andersson, 2017).

Integrating filtration as a pre-processing step for the isolation of microorganisms could provide good quality and integrity DNA from sediment samples that can be preserved sufficiently well and kept at ambient temperatures before DNA extraction and library construction for HTS-analyses. Most of the studies on applying pre-processing sediment samples by sequential membrane filtration focused on the quality assessment and efficiency of the extracted metagenomic DNA. Solomon et al. (2016) demonstrated that community DNA with minimal shearing was obtained from pre-processing marine sediment samples and performed PCR amplification of the 16S rDNA gene to confirm that the filtration method isolated high-quality DNA. A similar protocol was employed to process arctic sediment samples to characterize the bacterial community structure by 16S rDNA amplicon sequencing (Kachiprath et al., 2017). However, there is no comprehensive information on the potential biases of sequential membrane filtration on the retained microbial taxa compared to its non-processed counterpart, specifically whether sample pre-processing via sequential filtration compare to non-processed community profiles for quantitative measurements of freshwater microbial diversity and community structure.

Here, we examined if there would be method-driven differences between non- and pre-processed sediment samples (represented by the collection filter) by sequential membrane filtration for microbial community profiling through16s rDNA amplicon sequencing. Specifically, we evaluated the impact of pre-processing on the quality and quantity of extracted DNA, PCR amplicon, resulting HTS-reads, microbial diversity, and community composition with the non-processed sediment as the basis of comparison. Given the assumption that membrane filters of different size fractions (i.e., samples filtered from membranes of different pore sizes) retain different microbial biomass, we also assessed the difference in relative abundances, composition, and diversity of microbial taxa retained between each filter fractions. We provided the first comparison of the two approaches using 16s rDNA amplicon sequencing for sediment-associated microbial community profiling. Understanding the influence of pre-processing sediment samples for community analysis would be vital for conceptualizing appropriate study designs for sediment-associated microbial community profiling through molecular methods.

## MATERIALS AND METHODS

### Sediment Collection and Sample Pre-processing

The Trinity River is a large gravel-bed river impounded by the Trinity Dam (164 m a.b.l. and 3020 million m^3^ storage) and the smaller Lewiston Dam (28 m a.b.l. and 18 million m^3^ storage) in northern California, USA. It is under current dam operating guidelines with a mean annual flood of approximately 180 m^3^/s (Gaeuman et al., 2014). Sediment samples from three sites (i.e., sites A and C are from up-welling zones; site B from a down-welling zone) were collected approximately 10 cm below the submerged surface of selected gravel bars in the Trinity River assessed in the study of Serrana et al. (2020). The samples were stored in 50 ml sterile falcon tubes and immediately fixed with 99.5% molecular grade ethanol upon collection. The collected sediment samples were mainly composed of coarse sediments ranging from 1 to 5 mm in diameter, containing smaller sand grains and fine particulate mass. Pre-processing of sediment samples was done two to four hours after collection.

The experimental procedure of the sediment-associated microbial community profiling employed in this study is illustrated in **Figure 1**. Subsamples of ~600 mg each were aliquoted for sequential membrane filtration. The subsamples were resuspended in separate 50 ml solutions containing 0.22 μm filtered river water with Tween 20 (at a concentration of 1 ml l^-1^ v/v), agitated and mixed via a magnetic stirrer for 30 min. The resuspended subsamples were then filtered through a pre-filter with a 10 μm pore size (Nuclepore™ hydrophilic membrane filter paper; Whatman, Tokyo, Japan), followed by a mid-filter of 5 μm pore size (Mixed cellulose ester membrane filter; Merck Millipore, USA) and finally through a 0.22 μm collection filter (Cellulose mixed ester membrane filter; Merck Millipore, USA). The pre-processed samples were then kept in 2 ml microcentrifuge tubes, immediately fixed with 99.5% molecular grade ethanol. For non-processed sediments, triplicate subsamples of 200 mg were taken from the collected samples preserved in 50 ml Falcon tubes with 99.5% molecular grade ethanol.

**Figure 1.**
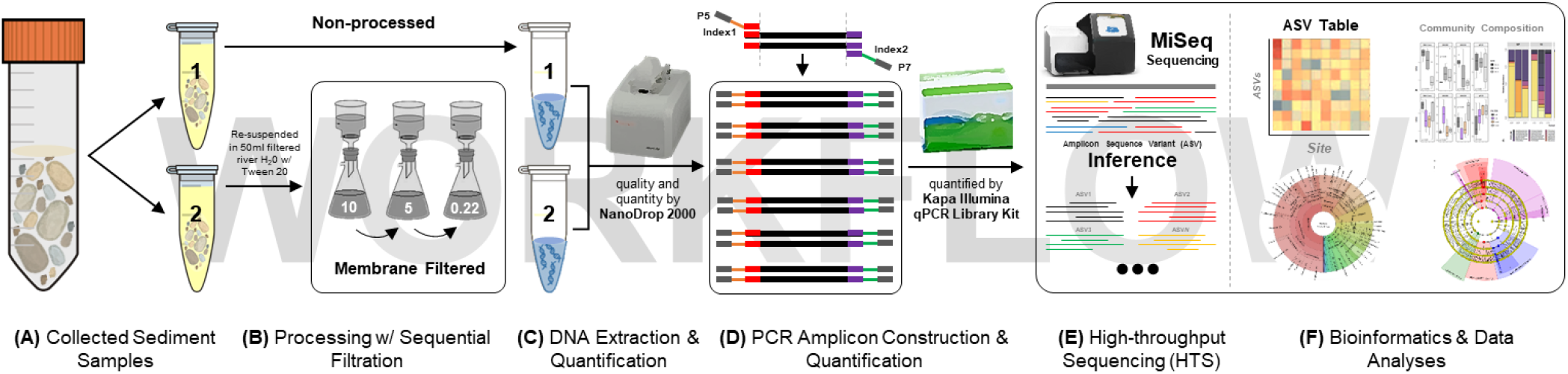
Schematic overview of the experimental procedure of the sediment-associated microbial community profiling employed in this study. Collection of sediment samples **(A)**. Sequential membrane filtration from 10, 5 to 0.22 μm pore size filters as pre-processing step **(B)**. DNA extraction following the protocol of Zhou et al. (1996) (as employed in Solomon et al., 2016) with some modifications **(C)**. One-step PCR amplification of the 16s rRNA V4 hypervariable region **(D)**. Sequencing through the Illumina MiSeq Platform **(E)**. Bioinformatics and statistical data analysis were done in R (R Core Team, 2019) **(F)**.

### DNA Extraction, PCR Amplification, and Sequencing

Before DNA extraction, the membrane filters were taken out from the collection tubes and dried at room temperature until most of the preserving ethanol evaporated. The membrane filter tubes (ethanol with finer particulate mass) and the subsampled non-processed sediments were then subjected to high speed (12,000 rpm) centrifugation for 30 min to resuspend the remaining fine particles and sediments to the bottom of each tube. The supernatant was removed carefully, and the tubes were dried at room temperature to evaporate the remaining ethanol. The dried membrane filters were cut into smaller pieces using sterile scissors and placed back into their original tubes. The samples were then suspended in a buffer consisting of 10 mM EDTA, 50 mM Tris-HCl, 50 mM Na_2_HPO_4_·H_2_O at pH 8.0 to remove PCR inhibitors (Zhou et al. 1996; Poulain et al., 2015). Genomic DNA was extracted from both the non-processed and filtered subsamples following the protocol of Zhou et al. (1996) (as employed in Solomon et al., 2016). The DNA extracted from non-processed sediment subsamples were combined accordingly before amplification. The quality and quantity of total DNA extracted was initially assessed with a NanoDrop spectrophotometer (NanoDrop 2000, Thermo Scientific).

Amplicon library preparation was carried out through a one-step PCR amplification using modified fusion primers of the V4 hypervariable region of the 16S SSU rRNA gene (i.e., 515F and 806R; Caporaso et al., 2012), with 12-base error-correcting Golay codes on both forward and reverse primers. The PCR was performed with high-fidelity Phusion polymerase (Thermo Fisher Scientific Inc.) in a T100 Thermal Cycler (Bio-Rad Laboratories, USA). The 25 μl PCR reaction mixture consisted of 5 μl of 5X Phusion GC Buffer, 1.25 μl each of the forward and reverse primers (10 μM), two μl dNTPs (2.5 mM), 0.75 μl DMSO, 0.25 μl Phusion Polymerase (1 U) and one μl of template DNA. The PCR condition followed was initial denaturation at 98°C for 3 min, 25 cycles of denaturation at 98°C for 15 s, annealing at 50°C for 30 s, and extension at 72°C for 30 s, followed by a final extension period at 72°C for 7 min.

Post-amplification, library-quality control was performed by checking the library size distribution via the High-Sensitivity DNA chip (Agilent BioAnalyzer). The libraries were purified and size selected using SPRI beads (AmpureXP, Beckman Coulter Genomics). Amplicon size was ~400-bp. Triplicate quantitative PCR reactions at appropriate dilutions were performed to quantify the amplicon libraries with the KAPPA Illumina Library qPCR Quantification kit (Kappa Biosystems, Wilmington, MA, USA). Negative control was used to monitor contamination from DNA extraction and PCR to post-amplification library quantity and quality verification; however, no quantifiable amplicon was detected for further analysis. The purified amplicon libraries were then normalized, and equimolar amounts were pooled. The 4 nM pooled library was sequenced at the Advanced Research Support Center (ADRES) of Ehime University using the Illumina MiSeq platform with paired-end reads of 300-bp per read.

### Read Processing and Taxonomic Assignment

The raw sequence reads generated on the Illumina MiSeq platform were demultiplexed via the command-line tool Cutadapt v.2.1 (Martin, 2011). The 3,805,575 demultiplexed sequences were quality screened, processed, and inferred amplicon sequence variants (ASVs) with the denoising pipeline of the DADA2 v.1.12 package (Callahan et al., 2016) in R v.3.6.2 (R Core Team, 2019). Based on the read error profiles, the reverse reads have poor read quality. Low read abundance with acceptable overlaps between the reads can be accounted for after quality filtering; therefore, only the forward reads were used in the subsequent analysis. Primer contaminants were excluded, and the reads were filtered based on quality and identified sequence variants likely to be derived from sequencing error. ASVs were inferred from the sequence data, subsequently removing chimeric sequences and singletons. The DADA2 pipeline was implemented to use sequence error models to correct amplicon errors in ASVs. Reads with a maximum expected error greater than 5 were discarded as a quality filtering measure and truncated at a read length of 100-bp. The remaining ASV sequences were aligned to the SILVA database (Pruesse et al., 2007) through the SILVA ACT: Alignment, Classification, and Tree Service online server (www.arb-silva.de/aligner) (Pruesse, Peplies, and Glöckner, 2012). For this analysis, the small subunit (SSU) category was selected, and a minimum similarity identity of 0.95 was set with ten neighbors per query sequence. Sequences below 70% identity were rejected and discarded. The least common ancestor (LCA) method was used for the taxonomic assignment. Chloroplasts, mitochondria, and unclassified ASVs were removed, resulting in a total of 2,875 taxonomically assigned ASVs.

The raw sequence data were deposited into the National Center for Biotechnology Information (NCBI) Sequence Read Archive (SRA) under the accession number PRJNA559761. The ASV matrix, the taxonomy, and sample table generated in this study have been deposited in the Figshare data repository (10.6084/m9.figshare.13088834).

### Statistical Analysis and Data Visualization

Statistical analyses were performed in R v.3.6.2 (R Core Team, 2019). The significant differences in the quality and quantity of extracted DNA and PCR amplicon libraries, and the HTS-reads for each read processing steps between sites (i.e., A, B, C), and filters [i.e., non-processed (NP), pre-filter (10 μm filter, “10”), mid-filter (5 μm, “5”), and the collection filter (0.22 μm, “0.22”)] were tested via two-way analysis of variance (ANOVA), and pairwise comparisons via multiple T-tests in the presence of significant main effects using the stat_compare_mean() in the ggpubr package (Kassambara, 2018). The correlation between the extracted DNA and PCR amplicon library concentration and purity, and between HTS-read count per processing step (i.e., raw reads, quality filtering, denoising, chimera removal, taxonomic assignment, and ASV count) were tested with Pearson correlation analyses on log-transformed data. A correlogram with significant tests was calculated and visualized with the Hmisc and corrplot packages (Harrell and Harrell, 2019).

Before subsequent statistical analyses, the ASV table was normalized at median sequencing depth. The shared and unique taxonomic assignment and ASVs between the groups were visualized with Venn diagrams and UpSetR plots (Lex et al., 2014). The boxplots were illustrated via ggplot2 (Wickham, Chang, and Wickham, 2016). The spatial differences between the microbial communities were visualized using non-metric dimensional scaling (NMDS) based on Bray-Curtis distances with the plot_ordination() function from the phyloseq package (McMurdie and Holmes, 2013), and in a hierarchical clustering dendrogram based on the average-linkage algorithm using the hclust() function. PERMANOVA (permutational multivariate analysis of variance) (vegan; Oksanen et al., 2013) was performed to identify significant differences in community composition between filters based on the NMDS ordination.

Alpha diversity metrics (i.e., Chao1 richness, Shannon diversity, Pielou’s J evenness, Berger-Parker’s dominance, and rarity index) were calculated and visualized based on the ASV dataset to identify the changes in community structure between the non-processed and filtered samples using the plot_alpha_diversities() function (microbiomeutilities; Sudarshan, Shetty and Lahti, 2018). Significant differences between the alpha diversity of sites and filters were also tested via ANOVA and pairwise comparisons via multiple t-tests in the presence of significant main effects. Linear discriminant analysis (LDA) effect size (LEfSe) (Segata et al., 2011) was performed using the Python LEfSe package (parameters: p < 0.05, q < 0.05, LDA > 2.0) to identify which microbial taxa significantly explained differences in community composition between the filter groups (i.e., NP, 10, 5, 0.22). The LEfSe algorithm was used to determine indicator taxa considering both the abundance and occurrence of a particular taxon.

## RESULTS

### DNA Yield, PCR Amplicon, and HTS-read Abundance

The initial concentration and the ratio of absorbance (at 260/280 and 260/230) to assess the purity of extracted DNA were measured via spectrophotometry (**Table 1**; **Supplementary Figure 1A** and **1B**). The DNA yield between sites (A, B, and C) and filters (NP, 10, 5, and 0.22) was higher for sites A and B, and NP and 0.22 filters, but a significant difference between the observed values were only reported for the sites. A ratio of ~1.8 is generally accepted as pure DNA for the 260/280 ratio. Although sites B and C, and filters 10 and 5 reported a relatively high 260/280 ratio, ANOVA showed no significant difference in DNA purity between sites and between filters. The 260/230 ratio was also relatively low for all samples given the accepted range of 2.0-2.2 for pure nucleic acid indicative of the presence of contaminants, e.g., EDTA, carbohydrates, and phenol. It was notable that the mean PCR amplicon library concentration of NP was relatively lower than those of the filtered samples, given that it has higher extracted DNA concentration. However, the PCR amplicon library concentrations quantified via qPCR were not significantly different between sites and between filters. The correlation between extracted DNA and PCR amplicon library concentrations was not significant (Pearson correlation: r = −0.024, p = 0.94) (**Supplementary Figure S2**).

**Table 1.** Quality and quantity of extracted DNA, PCR amplicon, and HTS-read and amplicon sequence variant (ASV) count per sediment sample.

| Site | Code | Filter Type | Extracted DNA<sup>a</sup> |  |  | Amplicon Library |  |  | Read Processing |  |  |  | Taxonomic Assignment |  |
| --- | --- | --- | --- | --- | --- | --- | --- | --- | --- | --- | --- | --- | --- | --- |
|  |  |  | ng/ul | A<sub>260</sub>/A<sub>280</sub> | A<sub>260</sub>/A<sub>230</sub> | qPCR (nM)<sup>b</sup> | Size (bp) | Band | Raw Reads | Quality Filtered | Denoiised | Non-Chimeric | Reads w/ Tax. ID | ASVs w/ Tax ID |
| A | ANP | NP | 307.1 | 1.4 | 0.46 | 4.88 | 410 | 1 | 1,166,991 | 560,891 | 456,284 | 446,586 | 424,397 | 461 |
|  | A02 | 0.22 | 479.7 | 1.37 | 0.61 | 14.16 | 409 | 2 | 305,045 | 171,106 | 154,579 | 147,490 | 143,019 | 436 |
|  | A05 | 5 | 110.1 | 1.43 | 5 | 45.97 | 408 | 1 | 73,962 | 40,433 | 32,181 | 31,458 | 30,146 | 183 |
|  | A10 | 10 | 619.2 | 1.45 | 0.37 | 56.53 | 413 | 1 | 260,341 | 148,596 | 112,796 | 105,220 | 96,643 | 633 |
| B | BNP | NP | 625.5 | 1.45 | 0.3 | 56.79 | 413 | 1 | 446,984 | 135,890 | 87,857 | 86,948 | 85,109 | 75 |
|  | B02 | 0.22 | 397.9 | 1.61 | 0.75 | 27.33 | 414 | 1 | 79,771 | 29,699 | 22,339 | 22,153 | 20,745 | 104 |
|  | B05 | 5 | 44.3 | 2.53 | 0.06 | 8.34 | 415 | 1 | 146,808 | 101,535 | 86,288 | 82,169 | 77,590 | 465 |
|  | B10 | 10 | 35.8 | 3.08 | 0.04 | 17.38 | 415 | 1 | 182,665 | 135,387 | 125,620 | 112,804 | 105,066 | 1,071 |
| C | CNP | NP | 107.1 | 1.67 | 0.13 | 1.44 | 396 | 1 | 790,386 | 381,687 | 285,719 | 275,483 | 113,715 | 426 |
|  | C02 | 0.22 | 2.3 | 1.86 | 0.09 | 76.9 | 412 | 2 | 112,123 | 61,001 | 54,375 | 50,476 | 46,631 | 295 |
|  | C05 | 5 | 5.1 | 1.03 | 0.17 | 43.23 | 411 | 3 | 23,323 | 14,327 | 12,515 | 10,757 | 9,582 | 141 |
|  | C10 | 10 | 7.6 | 5.1 | 0.03 | 4.4 | 388 | 2 | 217,176 | 138,449 | 97,116 | 94,199 | 89,877 | 460 |
| **Total** |  |  |  |  |  |  |  |  | **3,805,575** | **1,919,001** | **1,527,669** | **1,465,743** | **1,242,520** | **2,875** |

<sup>a</sup>Initial quantification and quality assessment of extracted DNA via NanoDrop Spectrophotometer. <sup>b</sup>Amplicon library quantification via Kappa Illumina Library Quantification Kit.
<sup>c</sup>DNA Assay for fragment size quantification and quality via Agilent 2100 BioAnalyzer High Sensitivity DNA Kit. "NP" stands for non-processed sediment samples; "10" for the pre-filter (10 µm), "5" for the mid-filter (5 µm), and "0.22" for the collection filter (0.22 µm).

Based on the site and filter grouping, sites A and C and filters NP, 10, and 0.22 had higher read abundances (from raw reads to reads with taxonomic assignment) and ASV counts than site B and filter 5, respectively (**Supplementary Figure S3** and **Supplementary Table 1**). ANOVA showed no significant difference in read and ASV counts between the sites, while the raw, filtered (ANOVA; p < 0.05), denoised, and non-chimeric reads (ANOVA; p < 0.10) were significantly different between the filters. Although the amplicon libraries were normalized to equimolar concentrations before HTS, the NP samples had significantly higher absolute raw read abundance than the filtered samples (t-test: p < 0.05). After quality filtering, NP was only significantly different against the 5 filters (t-test: p = 0.047). Furthermore, the correlations between the read abundances from raw reads to each processing step were all significantly (p < 0.05) positive with strong (Pearson’s r > 0.60) to very strong (Pearson’s r > 0.80) correlations (**Supplementary Figure S2**).

### ASV Richness, Taxonomic Diversity, and Community Composition

From the 2,875 ASVs, 2,871 were identified as bacteria, while 4 ASVs were assigned as archaea (i.e., Nitrosopumilales and Woesearchaeales). We identified a total of 324 microbial genera from 232 families under 161 orders, 85 classes, and 39 phyla, including unclassified taxa (e.g., Unclassified Bacteria). **Figure 2A** presents the relative abundance of the sediment-associated microbial phyla grouped per filter. Phyla with high relative sequence abundances include the Proteobacteria, Bacteroidota, and Acidobacteria (**Figure 2B).** Rhodobacteriaceae and Vicinamibacteriaceae predominantly represented non-processed sediments. Whereas Chitinophagaceae, Microscillaceae, and *Flavobacterium* dominate the 10, 5, and 0.22 filters, respectively (**Supplementary Figure S4**).

**Figure 2.**
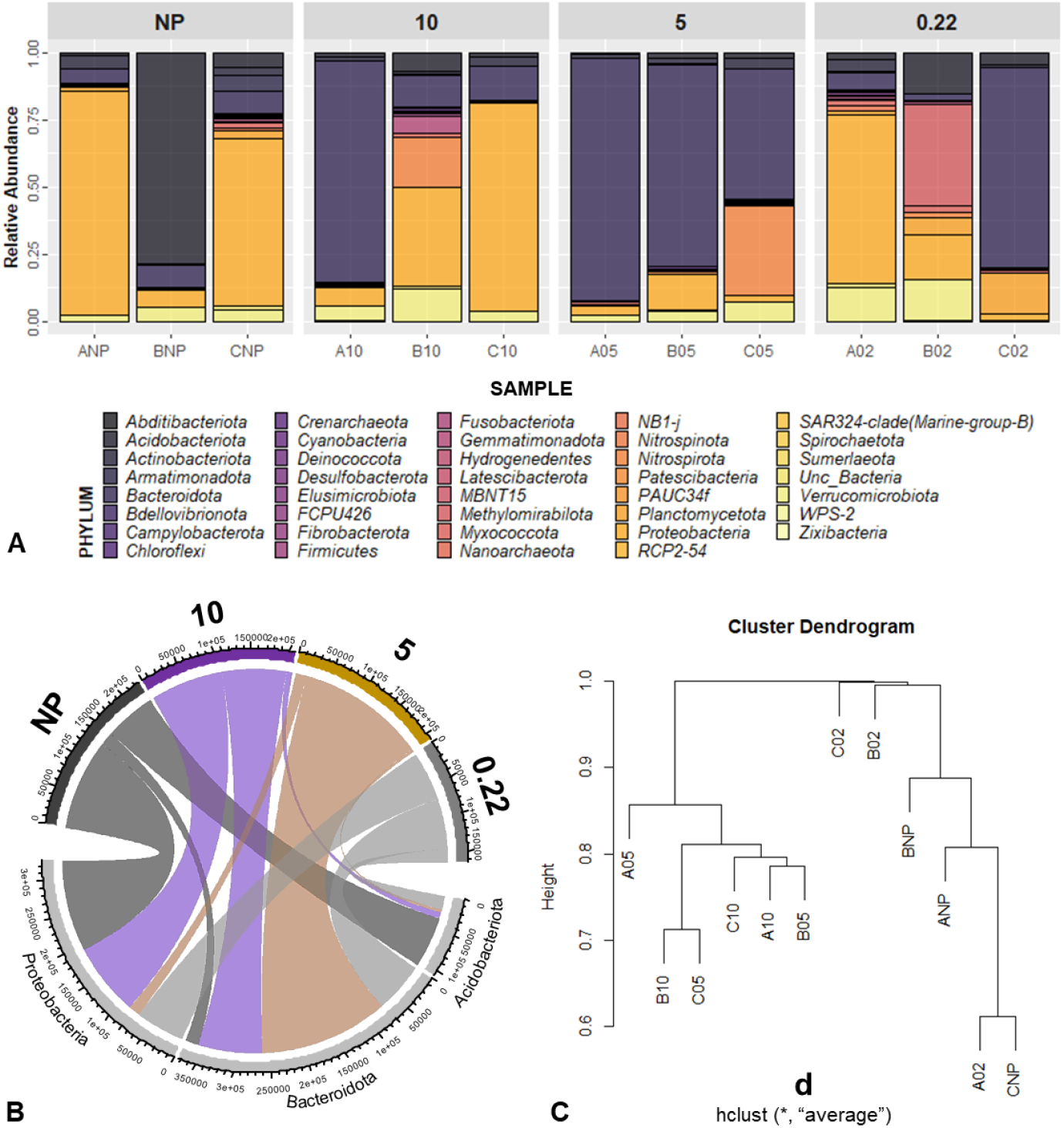
Relative abundance of microorganisms identified by 16s rDNA amplicon sequencing **(A)**. Compositions are illustrated at the phylum level. The chord diagram indicating the log-transformed abundance of the top three Phylum detected for each filters **(B)**. Hierarchical clustering dendrogram of the similarity in community composition across the sampling sites **(C)**.

To explore the difference between the non-processed and collection filter samples, the shared and unique ASVs and taxa (e.g., Phylum, Class, Order, Family, and Genus) assigned per filter were visualized via Venn diagrams (**Figure 3A** and **Supplementary Figure S5**) and UpSetR plots (**Figure 3B** and **Supplementary Figure S6**). Notably, the 10 filters always showed the highest ASV count throughout the sites (**Table 1**). When grouped by filter type, the 10 filters had the highest unique ASV count with 978, followed by 0.22, NP, and 5 with 594, 492, and 121 unique ASVs, respectively. The NP and 0.22 collection filters shared 63 + 89 (Mean + SD) or a total of 239 ASVs (74% of reads shared) having 257 + 143 (total of 493; 16% of reads) and 215 + 81 (total of 595; 10% of reads) unique ASVs, respectively.When aggregated at the genus level, the two methods shared 35 + 34 or a total of 108 genera (95% of reads) with 54 + 40 (total of 51; 2% of reads) and 39 + 1 (total of 59; 3% of reads) unique genera, respectively. Also, the 10 and 5 filters shared 449 ASVs, and no ASV was shared between all four filters.

**Figure 3.**
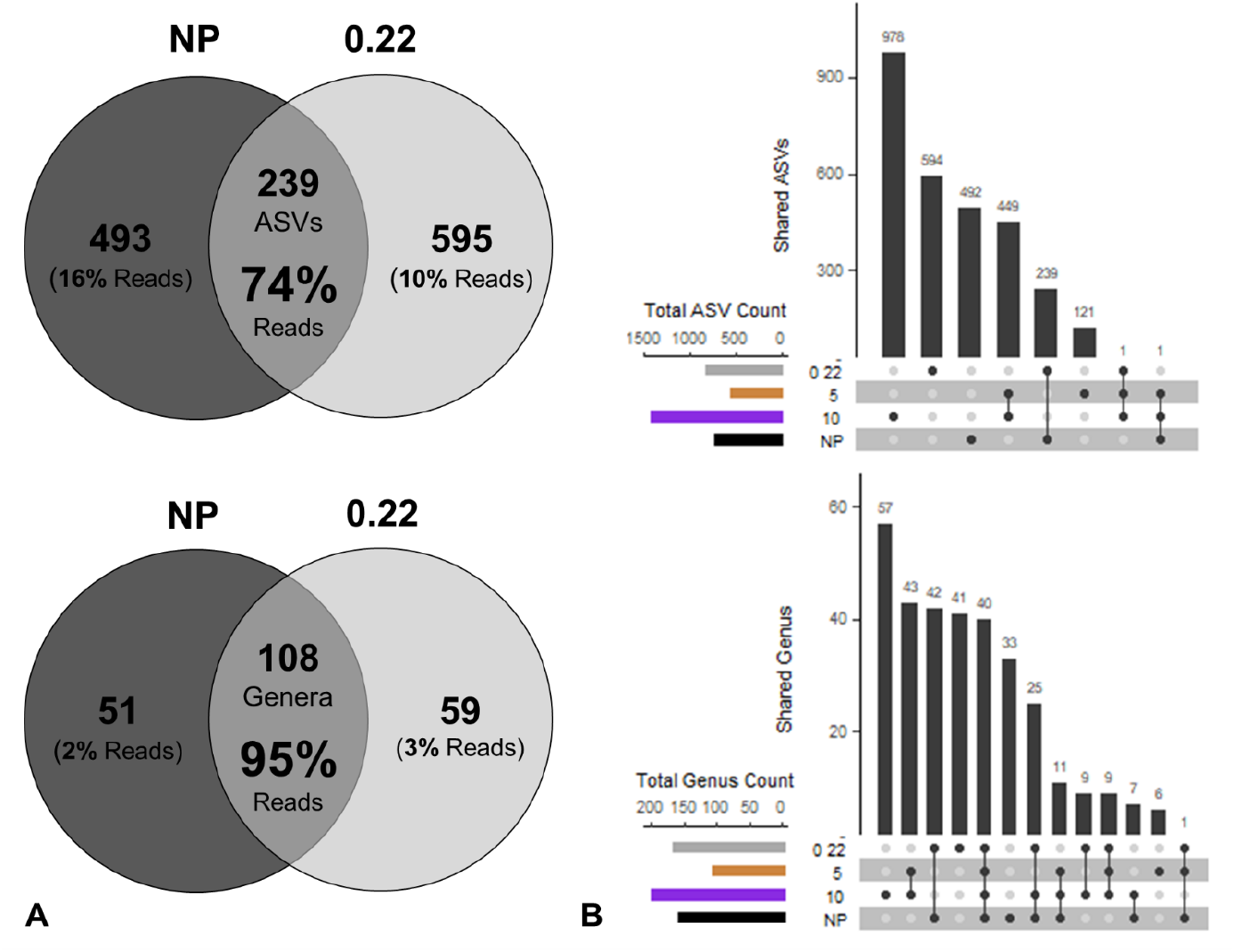
Venn Diagrams (A) and UpSetR plots (B) showing shared and unique ASVs and genus between the non-processed (NP) and pre-processed samples (represented by the collection filter, 0.22 μm), and between all groups (NP, 10, 5, and 0.22 μm) sediment samples. The bars in the upset plot show the overlap between the indicated sample below.

Alpha diversity based on Chao1 richness, Shannon diversity, Pielou’s evenness, Berger-Parker’s dominance, and the rarity index is presented in **Supplementary Figure S7**. ANOVA showed no significant difference between the sites and between filters in richness, diversity, evenness, dominance, and rarity estimates. Both the NMDS ordinations of the genus and ASV datasets indicated that the samples cluster based on the filters as visualized in the ordination space (**Supplementary Figure S8**). Notably, filters 10 and 5, and NP and 0.22 clustered closely together. The hierarchical clustering of samples based on the ASV dataset also showed the separation of NP and 0.22 against the 10 and 5 filters (**Figure 2C**). However, PERMANOVA showed no significant difference in the community composition of both the genus (R^2^ = 0.21, p = 0.245) and ASV (R^2^ = 0.22, p = 0.062) datasets.

### Indicator Taxa Analysis

LEfSe was performed to identify taxa significantly explained differences in the community compositions between the filter groups. Thirty-five significantly discriminative features out of 51 were selected before internal Wilcoxon, and 25 had an LDA score > 2. A cladogram showing the 25 microbial taxa’s phylogenetic distribution significantly associated with each filter group is presented in **Figure 4A**. The corresponding LDA values for each taxon are shown in **Figure 4B**.

**Figure 4.**
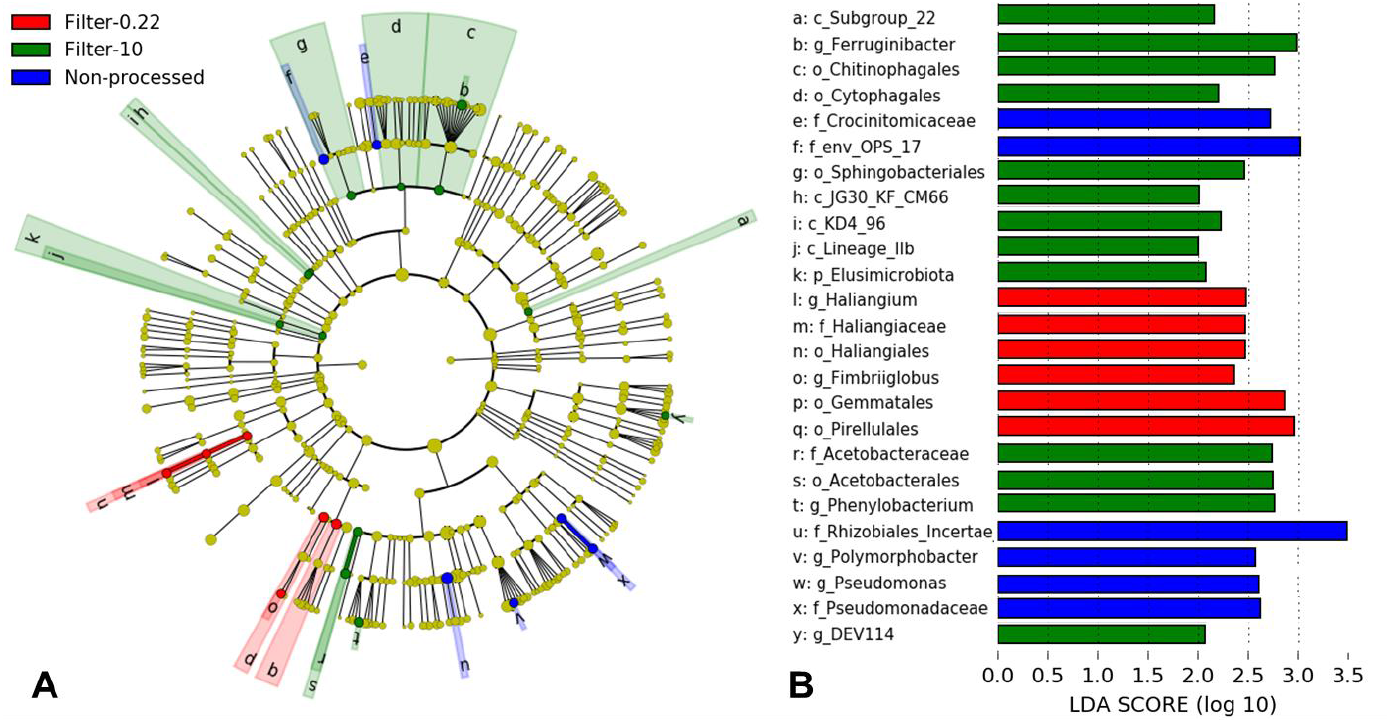
Linear Discriminant Analysis (LDA) Effect Size (LEfSe) plot of indicator taxa identified from non-processed (NP), and sequential filtered (10, 5, and 0.22 μm) sediment samples. Cladogram representing the hierarchical structure of the indicator taxa identified between the non-processed and filtered samples (filter) (A). Each filled circle represents one indicator taxa. Blue, indicator taxa statistically overrepresented in “NP”; red indicator taxa statistically overrepresented in “0.22”; green, indicator taxa statistically overrepresented in “10”. Identified indicator taxa grouped by filter and ranked by effect size (B). The threshold for LDA score was >2.0.

LEfSe analysis showed that the taxa from four families (i.e., Crocinitomicaceae, Env. OPS 17, Pseudomonadaceae, Rhizobiales *Incertae Sedis),* and two genera (i.e., *Polymorphobacter*, *Pseudomonas*) were significantly abundant in NP compared to other filter groups. For the sequential membrane filters, phylum Elusimicrobiota, four classes [e.g., Subgroup 22 (Acidobacteriota), JG30-KF-CM66 (Chloroflexi)], four orders (e.g., Chitinophagales, Sphingobacteriales), family Acetobacteraceae, and three genera [i.e., DEV114 (Pedosphaeracea), *Ferruginibacter, Phenylobacterium]* were significantly more abundant for the 10 μm filter, while three orders (i.e., Gemmatales, Haliangiales, Pirellulales), family Haliangiaceae), and two genera (i.e., *Haliangium, Fimbriiglobus)* were significantly more abundant for the 0.22 μm filter. No taxa were found to be significantly abundant for the 5 μm filter.

## DISCUSSION

In this study, we assessed whether riverine sediment-associated microorganisms would differ between non-processed and pre-processed samples by sequential membrane filtration. We provided the first comparison of the two approaches using 16s rDNA amplicon sequencing for microbial community profiling. We report that although the non- and pre-processed samples (represented by the final collection filter, 0.22) had more unique than shared ASVs, the latter accounted for a total of 239 ASVs that includes 74% of the reads between the two methods. More so at the genus level, the non- and pre-processed samples had a relatively high percentage of total shared genus count (108 genera, 50%) that accounts for 95% of the reads’ absolute abundance. This showed that the final collection filter (0.22) captured most of the abundant genus identified from the non-processed samples. Notably, the collection filter detected a total of 59 more unique genera (3% of the reads). These false-positive detections suggested that the pre-processed samples can detect taxa not captured from the non-processed approach.

A range of mechanisms potentially drove the false-positive detections. First, this could be due to the effectiveness of the multiple filtration process to reduce inhibitory compounds. Sequential-filter isolation techniques have been employed to improve the yield of environmental DNA by reducing the concentration of inhibitory compounds, e.g., humic acid, polysaccharides, metals, etc. (Solomon et al., 2016; Kachiprath et al., 2017; Hunter et al., 2019). Specifically, sediment samples contain high humic substances, which are the primary compounds co-extracted with DNA that inhibits enzymes (e.g., *Taq* polymerase) in PCR reactions (Matheson et al., 2010). The reduction of these inhibition compounds could have led to the generation of false-positive taxa in relation to the non-processed samples. However, we observed no significant difference in the quality of extracted DNA to support reduced inhibitory compounds’ influence on the false-positive detections. Other reasons, such as sequencing depth (the total number of usable reads from the sequencing machine), have been reported to influence the rate of false-positive detections in metabarcoding studies (Ficetola, Taberlet, and Coissac, 2016). Insufficient sequence depth could result in the non-detection of rare taxa. For example, singletons (single sequence detection, or an OTU/ASV only present in one sample) are usually considered erroneous sequences or artifacts and are usually removed for subsequent analysis. Increasing the sequencing depth might result in an increase in these reads’ abundances in the sample. Also, method-specific or unique taxa could result from having abundant taxa with polymorphisms (Laroche et al., 2017). On the other hand, setting a more stringent parameter for quality filtering could reduce the rate of detecting false positives (Ficetola et al., 2015; Serrana et al., 2018). Given that we employed a relatively lax read quality filtering parameter in this study, the false positive detection could result from low-quality passing reads.

On the other hand, the false-negative taxa (51 genera; 2% of the reads) absent from the collection filter could be microbial groups that passed through the 0.22 μm pore-sized filter. As previously reported by Maejima et al. (2018), isolated bacteria from lake water samples belonging to the Proteobacteria, Bacteroidetes, Firmicutes, and Actinobacteria passed through a 0.22 μm pore size filter. The filtered fractions from < 0.2 μm filtered samples that were usually considered “sterile” were found to still contain miniature cells, ultramicrobacteria (i.e., bacteria whose cell size are smaller than 0.1 μm^3^) and slender filamentous bacteria (e.g., Oligoflexia, Proteobacteria) overlooking a broad diversity of filterable agents (Wang et al., 2007; Nakai 2020). However, we observed that the false-negative taxa had very low read abundance, which could be due to smaller cell size leading to low DNA yield. This suggests that the microbial groups that possibly passed through the 0.22 μm pore-sized collection filter were mostly low abundant taxa. Nonetheless, we observed a low read abundance of these false-positive and negative detections. As demonstrated from the diversity and community composition analyses employed in this study, these method-specific taxa would unlikely affect these results.

On the other hand, the pre- and mid-filters had a relatively high count of 449 shared and 978 and 121 unique ASVs, respectively. The non-processed samples only had 1 ASV shared with the pre- and mid-filter, similar to the collection filter. The clear separation between NP and 0.22 against the 10 and 5 filters was also observed in the NMDS ordination and the hierarchical clustering. At the genus-level, the pre- and mid-filters had 57 and 6 unique genera. These values added with the genera shared between the two filters makes a total of 106 captured solely from the pre- and mid-inline filtration. The very low ASV and low genera shared between non-processed and collection filter against the pre- and mid-filters suggested that a huge part of the sediment microbial community is underrepresented or lost from the community profile. A previous study comparing the prokaryotic and eukaryotic diversity and community composition between pre- and collection filters from lake water samples suggested the possible “pre-filter” bias in the community structure from the collected biomass (Lanzen et al., 2013). They reported contrasting read abundance even though most operational taxonomic units (OTUs) were shared between filters. Sequential filtration of sediments might be a stochastic process where taxa are presumably retained according to cell size rather than their abundance, with the rare taxa retained along the previous filtration step (Pinto et al., 2020). We presented a stronger pre- and mid-filter community composition bias, given that very few ASVs and taxa were shared between the in-line filters against the non- and pre-processed samples. Since we observed that certain sediment-associated microbial taxa were not captured from the non-processed samples, and if only the collection filter is considered to represent the pre-processed samples’ microbial community profile, we suggest the inclusion of pre-filters in microbial communities’ profiling.

Statistical analyses revealed that groups based on filter were not significantly different in the richness, diversity, and evenness estimates of alpha diversity. Although shared taxa between the two methods were relatively low, community structures based on Bray-Curtis distance were also not significantly different between the two methods. Bray-Curtis dissimilarity is sensitive to differences in abundance between taxa, where abundant taxa are weighted more than the rare ones (Ricotta and Podani, 2017). Although the overall microbial community composition was not significantly different between the two methods, the significantly abundant indicator taxa detected between the filter types were different, primarily due to the variations in the detection of low abundance taxa. Based on LEfSe, representatives from the Alphaproteobacteria (i.e., Rhizobiales *Incertae Sedis,* and *Polymorphobacter), Pseudomonas* (Pseudomonadaceae) and the Crocinitomicaceae and the uncultured eubacterium env. OPS 17 were significantly more abundant in the non-processed sediment samples. The taxa affiliated with the Alphaproteobacteria have shown a consistent preference for a particle-attached lifestyle (Mestre et al., 2018). The pre-filter (10 μm filter) had the most significantly more abundant taxa with representatives from Acetobacteraceae (Alphaproteobacteria), Acidobacteriota, Bacteroidota, Chloroflexi, and Elusimicrobiota. Candidate microbial divisions and Chloroflexi have been reported to be primarily recovered when particle samples were subjected to filtration in situ (Torres-Beltrán et al., 2019). The collection filter had significantly more abundant *Fimbriiglobus* (Gemmatales), Pirellulales, and *Haliangium* (Haliangiales) sequences. The first two taxa are classified as members of the Planctomycetes, while the latter belongs to the Myxococcota. A study evaluating the influence of standard filtration practices on marine particles also reported that proportional abundances in the pre-filter fraction of Myxococcales (Deltaproteobacteria) and Planctomycetes increased with filter volume (Padilla et al., 2015). Furthermore, in-situ filtration (0.4 μm filter) increased the capture of Planctomycetes by fivefold compared to on-ship in-line filtration (Torres-Beltrán et al., 2019).

The isolation and capture of good quality and quantity DNA from sediment samples are very challenging (Harnpicharnchai et al., 2007; Solomon et al., 2016), and the preservation medium and the time between collection and storage is critical for particle or sediment-associated microorganisms to prevent biased overgrowth and DNA damage before HTS sample processing (Song et al., 2016). We observed that extracted DNA concentration varied between sites and filters and was relatively high for the NP filters. However, no significant difference was observed for the DNA yield between the two methods. PCR amplicon concentration and quality were also not significantly different between the non-processed and processed samples. Hence, we report that the quantity and quality of extracted DNA and its PCR amplicon libraries were not significantly different between the non-processed and processed samples. We should note that we used the same DNA extraction method for both non-processed and processed samples, employing the method of Zhou et al. (1996), which includes the removal of PCR inhibitors, i.e., humic compounds. The chosen DNA extraction method could present different impacts on the characterization of the overall microbial community composition (Ushio, 2019). Previous studies have investigated the influence of filter types and pore sizes on DNA yield from aquatic ecosystems (i.e., on environmental DNA, e.g., Robson et al., 2016; Li et al., 2018). Filters of different pore sizes did not affect the amount of total DNA recovered and detected species from environmental DNA (Li et al., 2018).

The PCR amplicon libraries were normalized before sequencing to assure an even read distribution for all samples. However, the raw HTS-reads and quality-filtered reads were significantly different between methods, with the non-processed significantly having the highest raw read abundance. Interestingly, after the denoising and the chimeric-read filtering steps, the retained reads from the non-processed sample declined and were not significantly different between methods. This suggested that the retained read abundance after the bioinformatics step was not significantly influenced by sediment processing or lack thereof. Previous studies have reported that higher GC content and larger insert size decreased the abundance of reads retained after quality filtering (Huptas et al., 2016). Moreover, fragment length may also impact the base qualities of Illumina reads (Tan et al., 2019). The decline in read abundance of NP (from being significantly different from the others to insignificant difference) after quality filtering suggests the possibility of the extracted DNA having either high GC content or large fragments which reduced the base qualities of the reads.

Our time from collection to processing and ethanol preservation of the filtered samples were from two to four hours. Previous studies reported that larger processing time between sample collection and filter storage might allow the growth of opportunistic prokaryotic groups introducing bias by microbial population turnover within the sample (Puigcorbé et al., 2020). Here the sediments processed for sequential membrane filtration were from samples that have already been preserved in ethanol; hence, this bias was not tested in our experimental design. We recommend further assessment of sediment pre-processing by comparing different filter types and combinations, preservation medium, sample volume, and the influence of various processing time for further method evaluation. This will fully present the capability and viability of on-site sequential membrane filtration as a processing step against the direct collection and preservation of riverine sediment samples.

## CONCLUSION

In the present study, we found no significant difference in the quantity and quality of extracted DNA and PCR amplicon between non- and pre-processed sediment samples. Raw and quality-filtered reads were significantly different between methods, but read abundance after bioinformatics processing were not significantly different. These results suggest that read abundance after the bioinformatics steps were not significantly influenced by sediment processing or lack thereof. We report that although the non- and pre-processed sediment samples had more unique than shared ASVs, both methods shared a total of 239 ASVs that accounts for 74% of the reads. More so at the genus level, the final collection filter also detected most of the genus identified from the non-processed samples, with 51 false-negatives (2% of the reads) and 59 false-positive genera (3% of the reads). All of the alpha diversity indices estimated, and the microbial community composition was not significantly different between the non- and pre-xprocessed samples. These results demonstrate that while differences in shared and unique ASVs and microbial taxa were detected, both methods revealed comparable microbial diversity and community composition. We also suggest the inclusion of sequential filters (i.e., pre- and mid-filters) in the community profiling, given the additional taxa not detected from the non-processed and the final collection filter. We presented the feasibility of pre-processing sediment samples for community analysis and the need for further assessment sampling strategies to help conceptualize appropriate study designs for sediment-associated microbial community profiling.

## Author Contributions

JMS performed field sampling and sample processing. JMS and KW conceptualized the study, analyzed the data, and wrote the manuscript.

## Acknowledgments

We are grateful to Dr. Bin Li of the Molecular Ecology and Health Laboratory (MEcoH), Ehime University, members of the Disaster Prevention Research Institute (DPRI), Kyoto University, and Dr. David Gaeuman of the Trinity River Restoration Program (TRRP) for their assistance during the field survey. We thank Dr. Naohito Tokunaga of the Division of Analytical Bio-Medicine for his assistance in performing high-throughput sequencing on the Illumina MiSeq platform of the Advanced Research Support Center (ADRES), Ehime University.

## Supplementary Material

The Supplementary Material for this article is submitted as an attachment: MS-Sediment-Filtering_SuppMat.pdf.

## Data Availability Statement

The raw sequence data were deposited into the National Center for Biotechnology Information (NCBI) Sequence Read Archive (SRA) under the accession number PRJNA559761. The ASV matrix, the taxonomy and sample table generated in this study have been deposited in the Figshare data repository (10.6084/m9.figshare.13088834).

