## Supplementary Material for "Sediment-associated microbial community profiling: sample pre-processing through sequential membrane filtration for 16s rDNA amplicon sequencing"

Running Title: Pre-processing for Sediment Microbial Community Profiling

**Authors and Affiliations**

Joeselle M. Serrana[1,2,3] and Kozo Watanabe[1,2,3*]

[1]Center for Marine Environmental Studies, Ehime University, Bunkyo-cho 3, Matsuyama, Ehime 790-8577, Japan

[2]Graduate School of Science and Engineering, Ehime University, Bunkyo-cho 3, Matsuyama, Ehime 790-8577, Japan

[3]Biological Control Research Unit, Center for Natural Sciences and Environmental Research, De La Salle University, 2401 Taft Avenue, Manila 1004, Philippines

***corresponding author**

Prof. Kozo Watanabe, PhD

Phone & Fax Number: +81 (0) 89 927 9847

### Supplementary Data

The raw sequence data were deposited into the National Center for Biotechnology Information (NCBI) Sequence Read Archive (SRA) under the accession number PRJNA559761. The ASV matrix, the taxonomy and sample table generated in this study, and the scripts used to perform all bioinformatics analyses have been deposited in the Figshare data repository (10.6084/m9.figshare.13088834).

### Supplementary Figures

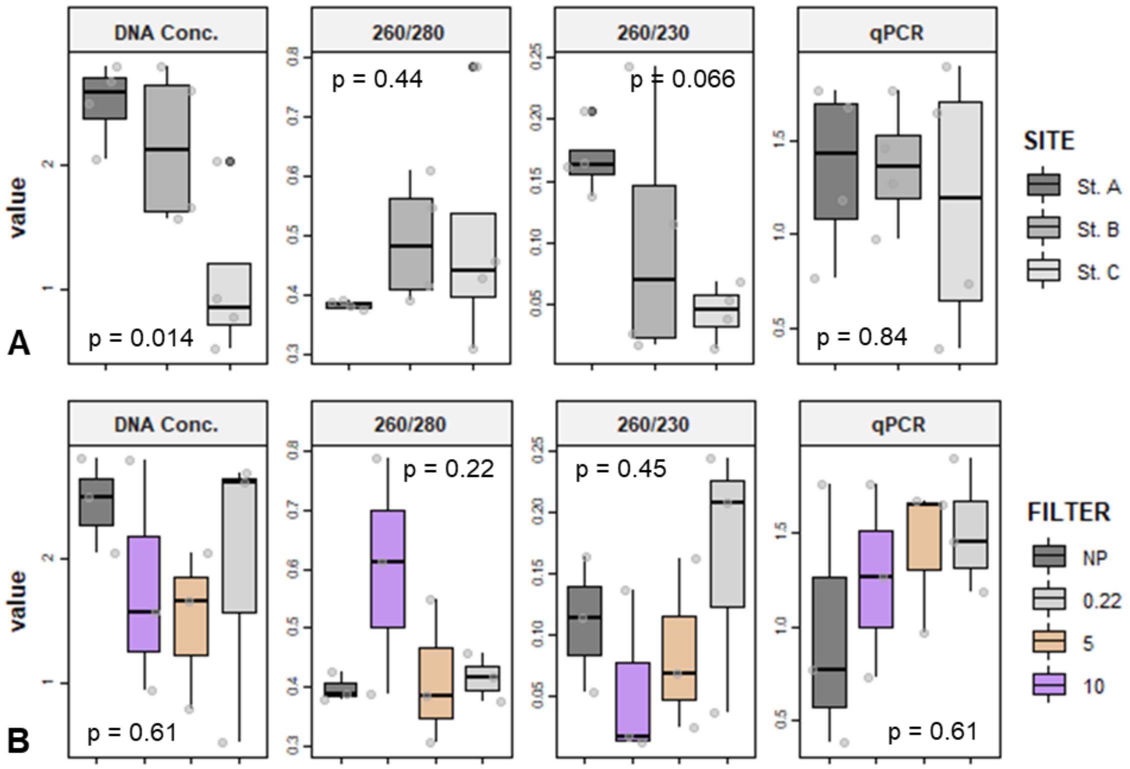


**Supplementary Figure 1.** Box-and-whisker plots of extracted DNA concentration and quality (log-transformed) grouped by site **(A)** and filter **(B)**. The p-value presented were from the analysis of variance (ANOVA) tests between samples.

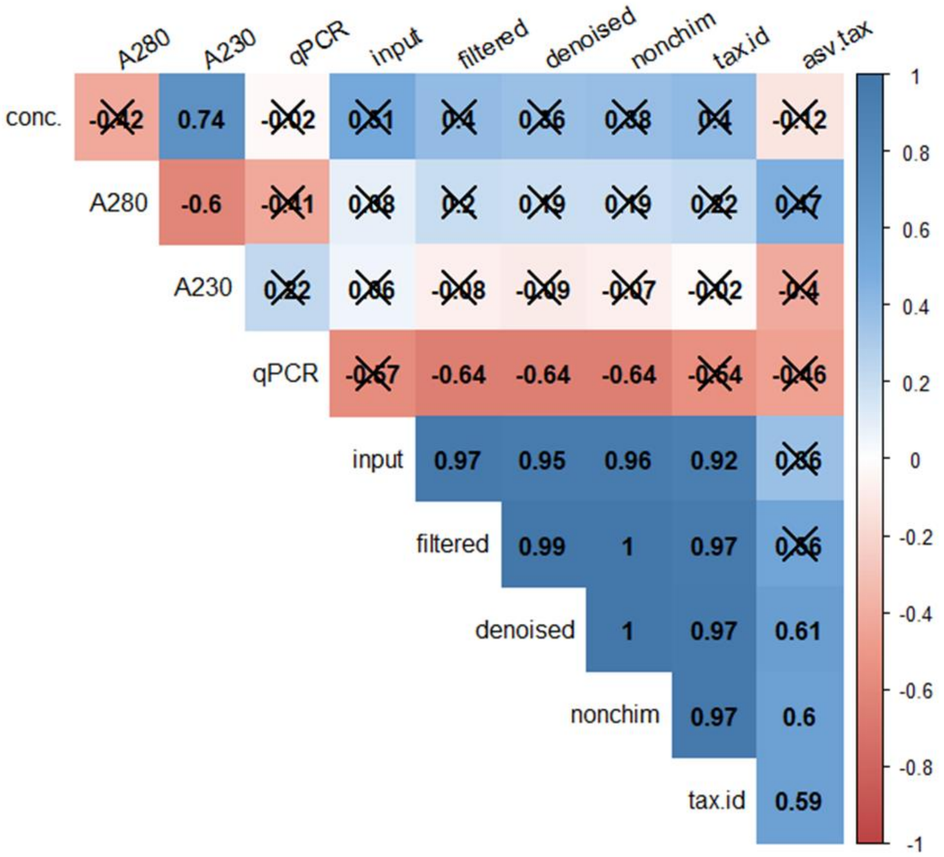


**Supplementary Figure 2.** Pearson's correlation matrix on log-transformed values. Statically significant ($p < 0.05$) Pearson's R values are highlighted. "conc." Stands for extracted DNA (ng/µl); "A280" for 260/280 ratio of DNA purity; "A230" for 230/280 ratio of nucleic acid purity; "qPCR" for the PCR amplicon library concentration (nM); "input" for the raw HTS-reads; "filtered" for quality filtered reads; "denoised" for the denoised reads; "nonchim" for the non-chimeric reads; "tax.id" for the reads with taxonomic assignment; "asv.tax" for the ASV count with taxonomic assignment.

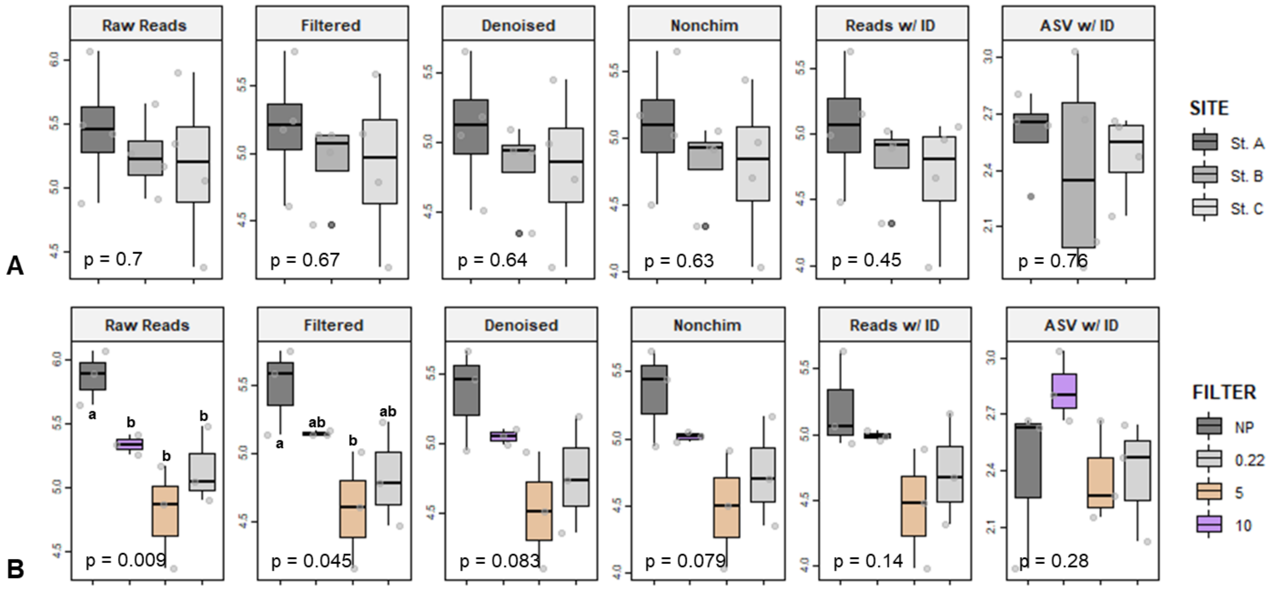


**Supplementary Figure 3.** Box-and-whisker plots of log-transformed read and amplicon sequence variant (ASV) abundance grouped by site **(A)** and filter **(B)**. Means with the same letter are not significantly different according to t-test at $p < 0.05$.

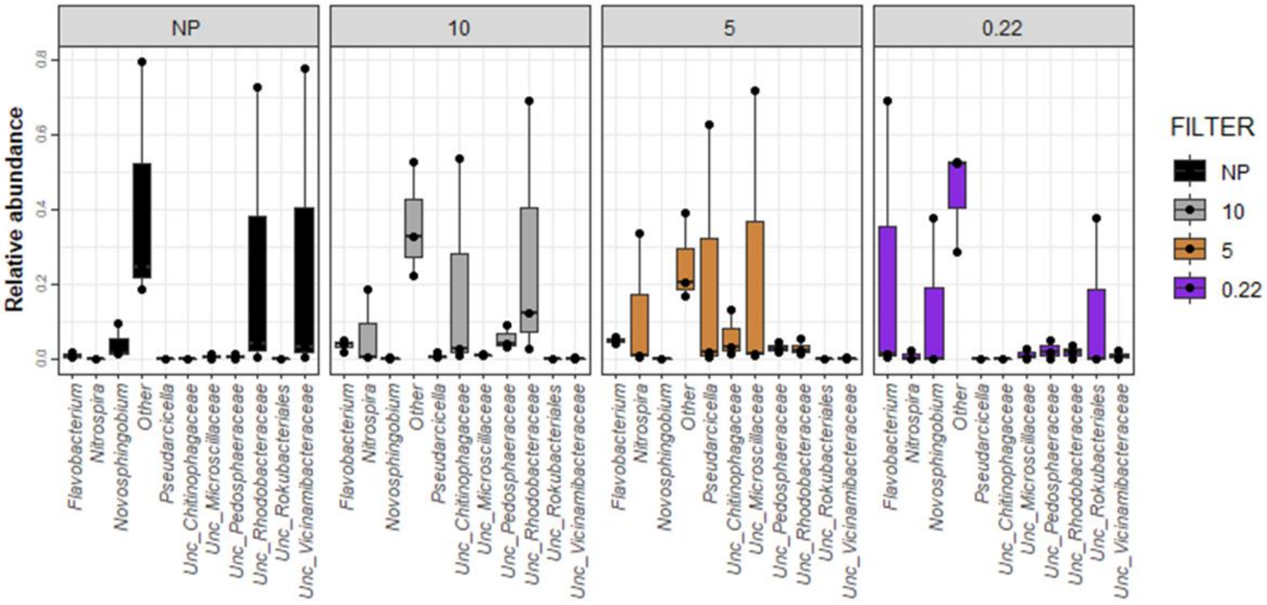


**Supplementary Figure 4.** Relative abundance of the top 10 genera grouped by filter.

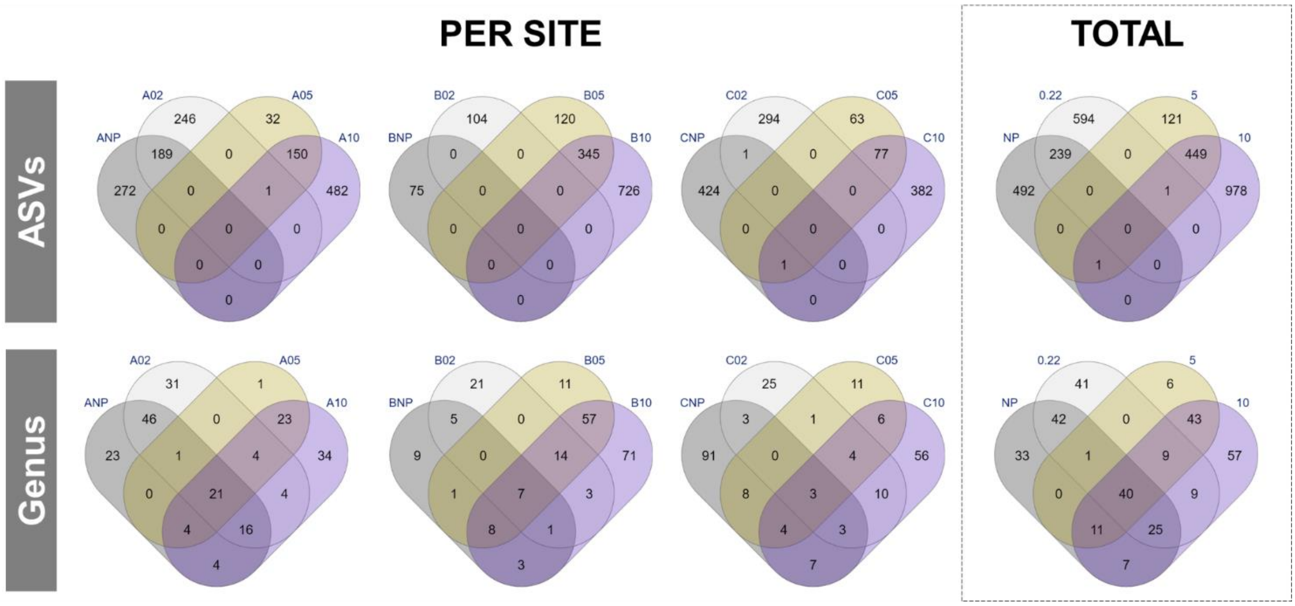


**Supplementary Figure 5.** Venn Diagrams showing shared and unique ASVs and genus between the filter types amongst sites.

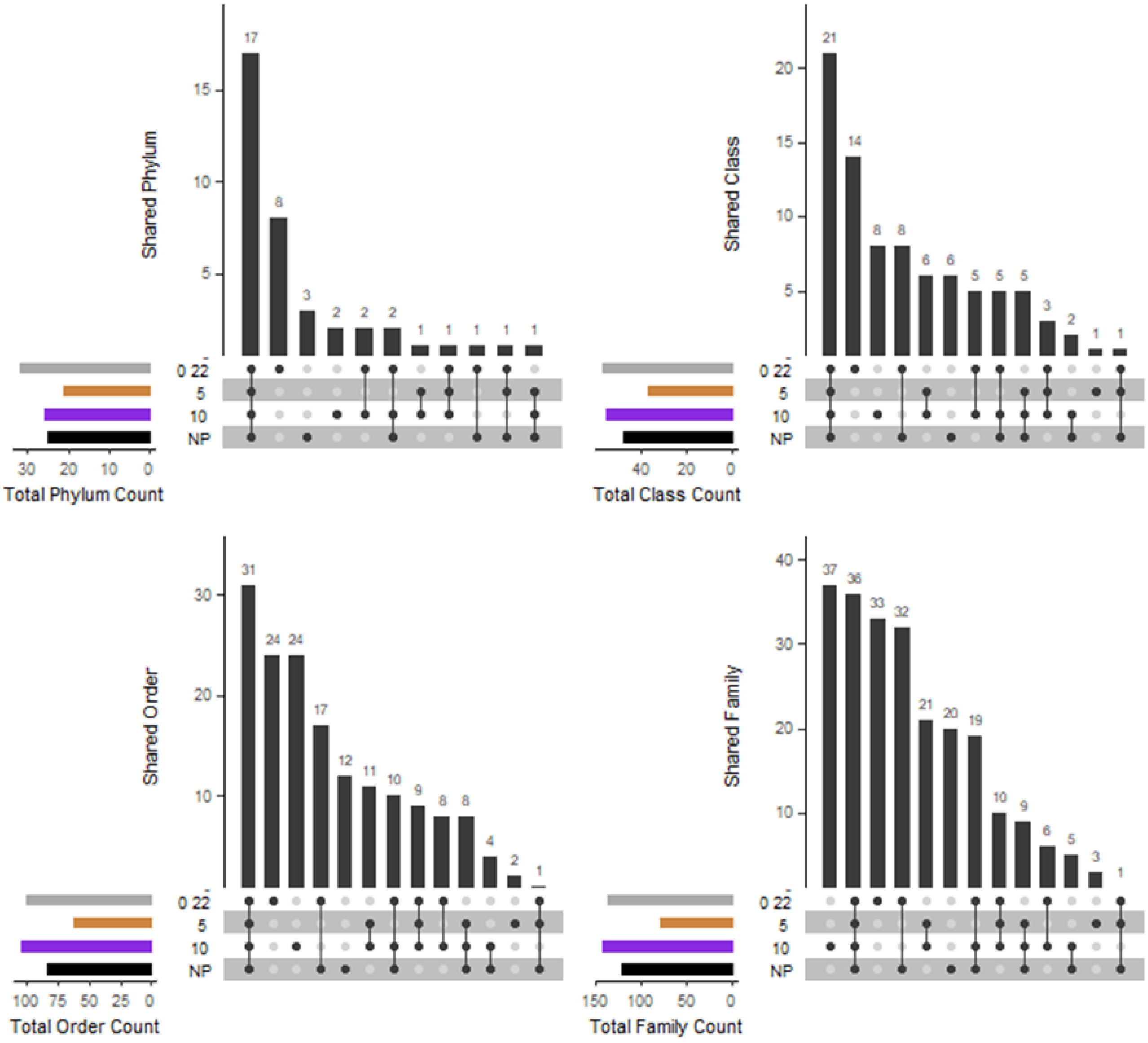


**Supplementary Figure 6.** UpSetR plots showing shared and unique taxa () between the non-processed (NP) and filtered (10, 5, and 0.22 µm) sediment samples. The bars in the upset plot show the overlap between the indicated sample below.

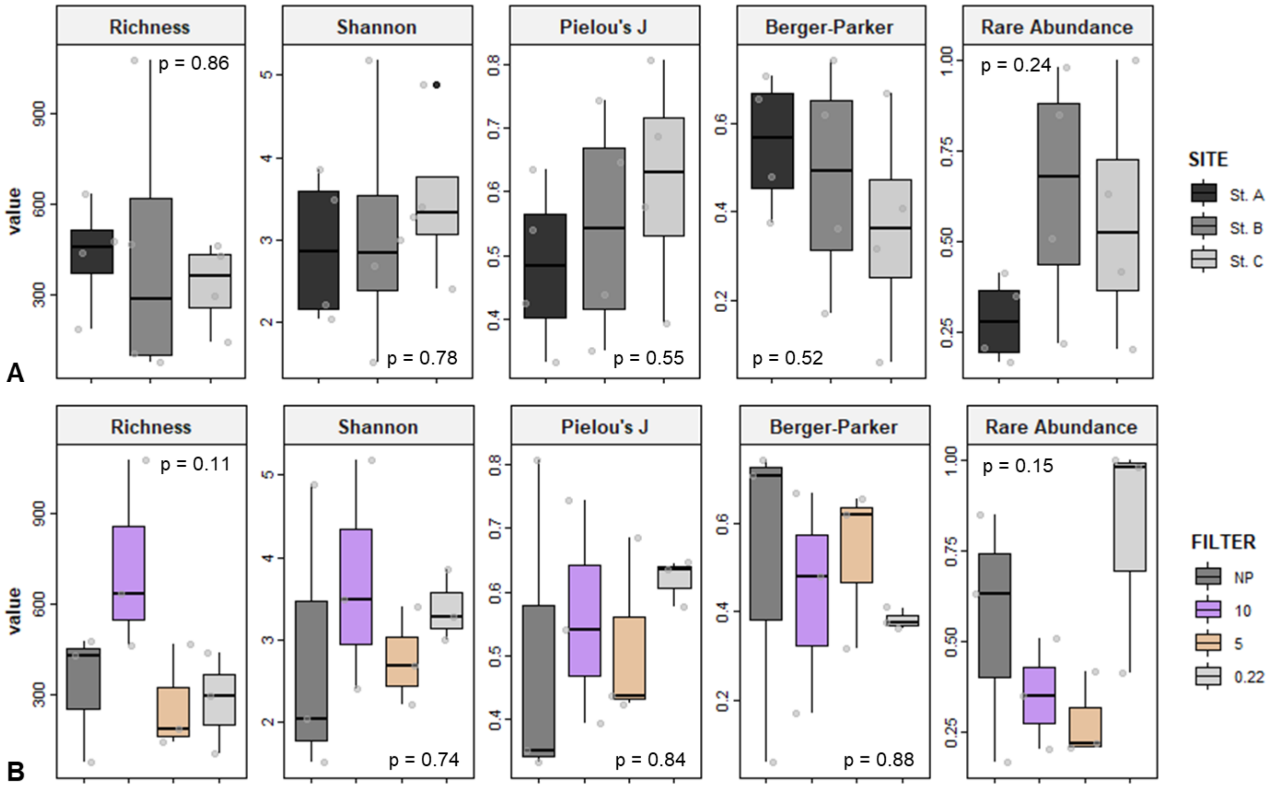


**Supplementary Figure 7.** Box-and-whisker plots of alpha diversity indices metrics comparing the samples by site **(A)**, and filter **(B)**.

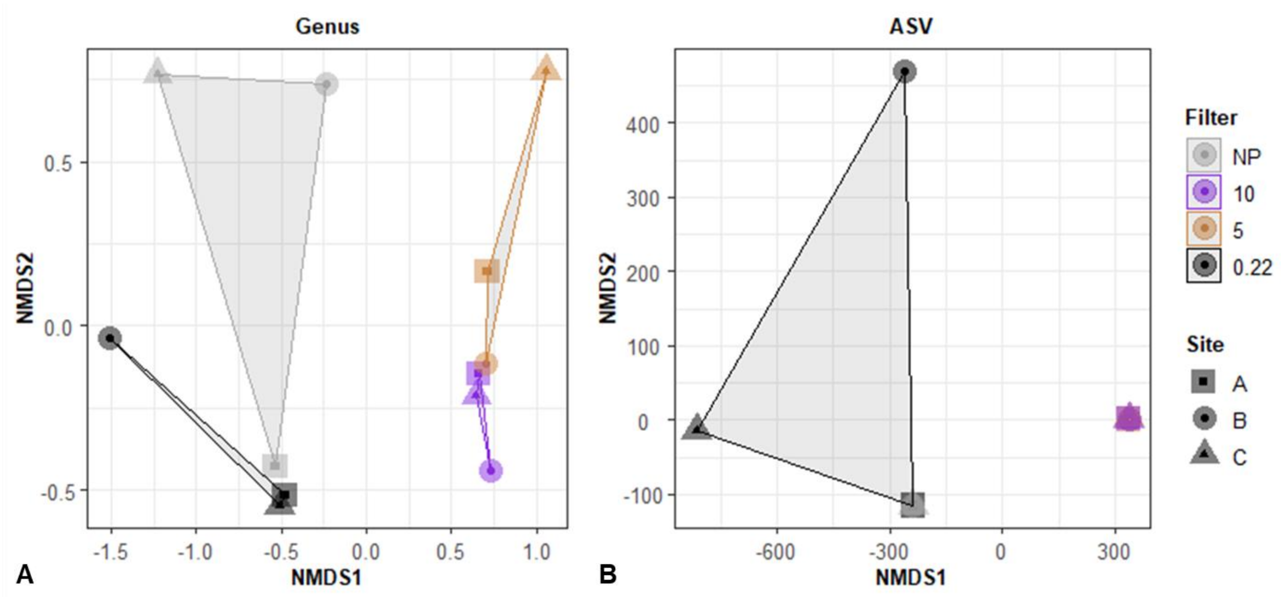


**Supplementary Figure 8.** Cluster analysis via non-metric multidimensional scaling (NMDS) based on Bray-Curtis dissimilarity showing microbial community composition the non-processed (NP), and sequential filtered (10, 5, and 0.22 μm) sediment samples for the genus **(A)** and ASV **(B)** datasets.

### Supplementary Table

**Supplementary Table 1.** Read and amplicon sequence variant (ASV) abundance grouped by filter [mean average (standard deviation)].

| Filter Type | Raw Reads | Quality Filtered | Denoised | Non-Chimeric | Reads w/ Tax. ID | ASVs w/ Tax ID |
|---|---|---|---|---|---|---|
| NP | 801,453 (360,131)$^{a}$ | 359,489 (213,368)$^{a}$ | 276,620 (184,381)$^{a}$ | 269,672 (179,889)$^{a}$ | 207,740 (188,174)$^{a}$ | 320 (213)$^{a}$ |
| 0.22 | 165,646 (121,801)$^{b}$ | 87,268 (74,272)$^{ab}$ | 77,097 (68,986)$^{a}$ | 73,373 (65,730)$^{a}$ | 70,131 (64,435)$^{a}$ | 278 (166)$^{a}$ |
| 5 | 81,364 (62,074)$^{b}$ | 52,098 (44,759)$^{b}$ | 43,661 (38,202)$^{a}$ | 41,461 (36,741)$^{a}$ | 39,106 (34,878)$^{a}$ | 263 (176)$^{a}$ |
| 10 | 220,060 (38,918)$^{b}$ | 140,810 (6,913)$^{ab}$ | 111,844 (14,275)$^{a}$ | 104,074 (9,355)$^{a}$ | 97,195 (7,609)$^{a}$ | 721 (314)$^{a}$ |

"NP" stands for non-processed sediment samples; "10" for the pre-filter (10 μm), "5" for the mid-filter (5 μm), and "0.22" for the collection filter (0.22 μm). Means with the same letter are not significantly different according to t-test at $p < 0.05$ (also shown in a boxplot in Supplementary Figure 3B).
